# Geometry alone influences stem cell differentiation in a precision 3D printed stem cell niche

**DOI:** 10.1101/489773

**Authors:** Elisabetta Prina, Laura Sidney, Maximilian Tromayer, Jonathan Moore, Robert Liska, Marina Bertolin, Stefano Ferrari, Andrew Hopkinson, Harminder Dua, Jing Yang, Ricky Wildman, Felicity Rose

## Abstract

Stem cells within epithelial tissues reside in anatomical structures known as crypts that are known to contribute to the mechanical and chemical milieu important for function. To date, epithelial stem cell therapies have largely ignored the niche and focussed solely on the cell population to be transplanted. Our aim was to recreate the precise geometry of the epithelial stem cell niche using two photon polymerisation and to determine the influence of this structure alone on stem cell phenotype. We were able to recreate crypt structures and following cell seeding, a zonation in cell phenotype along the z-axis emerged. This illustrates that geometry alone, without the use of exogenous signalling molecules, influences cell response. Understanding the role of geometry in the regulation of the stem cell niche will enable significant advances in our ability to influence stem cell behaviour to expedite cellular therapies to the clinic.

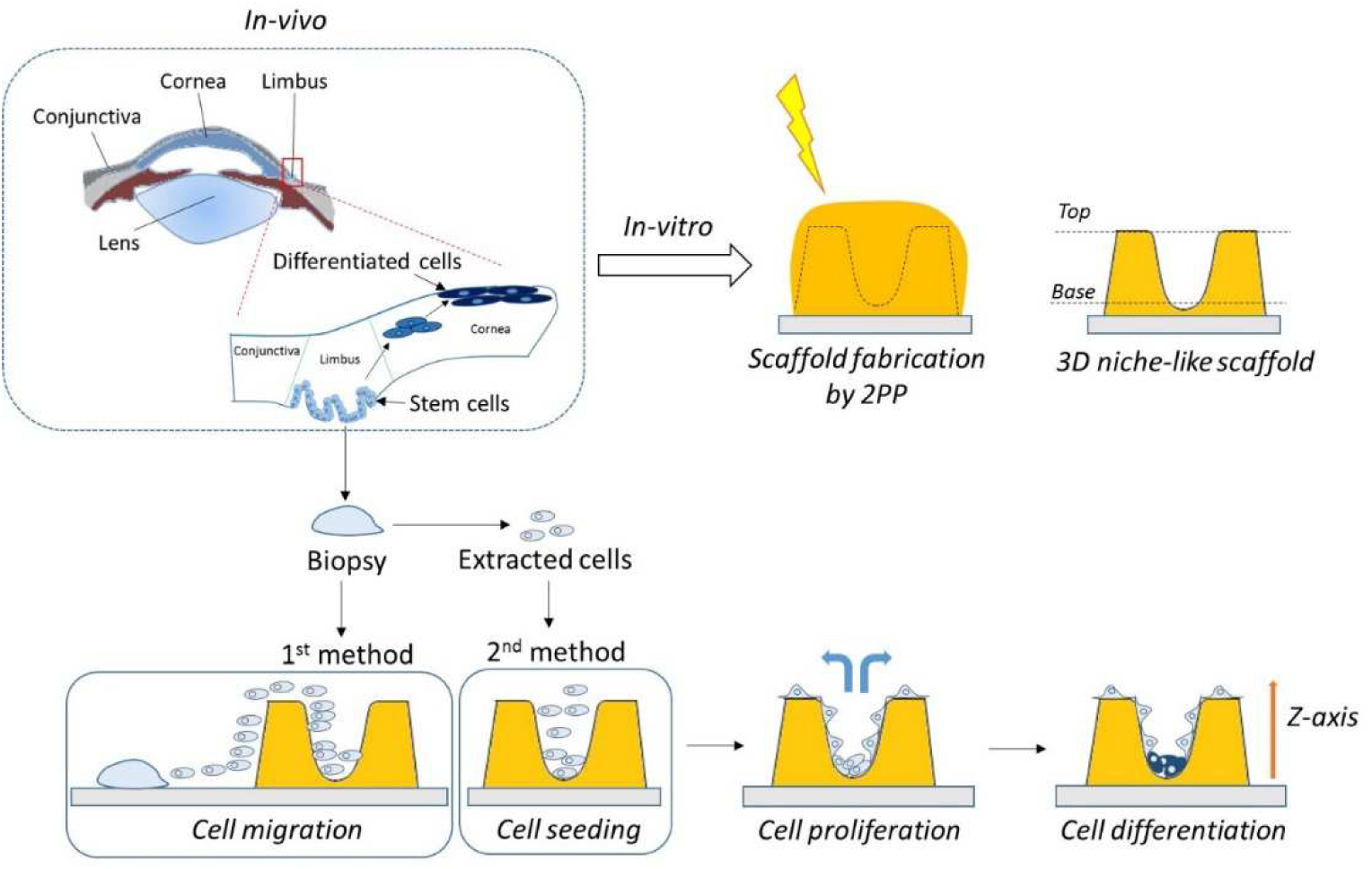
Graphical Abstract.

## Introduction

Stem cells within epithelial tissues reside in anatomical structures known as ‘crypts’ that are known to contribute to the mechanical and chemical milieu important for function. To date, epithelial stem cell therapies have largely ignored the niche and focussed solely on the cell population to be transplanted. Our aim was to recreate the precise geometry of the epithelial stem cell niche using two photon polymerisation and to determine the influence of this structure alone on stem cell phenotype. We were able to recreate crypt structures and following cell seeding, a zonation in cell phenotype along the z-axis emerged. This illustrates that geometry alone, without the use of exogenous signalling molecules, influences cell response. Understanding the role of geometry in the regulation of the stem cell niche will enable significant advances in our ability to influence stem cell behaviour to expedite cellular therapies to the clinic.

In the body, tissue maintenance and regeneration involves stem cell division, proliferation and differentiation. This process is tightly regulated in healthy tissue and is driven by a combination of intrinsic and extrinsic factors. Typical intrinsic elements include inherited polarity and nuclear factors controlling gene expression and chromosomal modifications^1^. Extrinsic factors include all the signals related to the stem cell microenvironment^2,3^. Stem cells are located in unique microenvironments, called niches, where they can reside for an indefinite period of time and produce progeny cells whilst self-renewing^4^. A disease state arises if changes in these factors lead to aberrant control or destruction of the stem cell niche. In regenerative medicine, a lack of control of stem cell behaviour is likely to contribute to high failure rates of stem cell therapies^5^. Current research in supporting epithelial regeneration has mainly focused on studying the effect of extracellular matrix composition, the cell-to-cell interactions mediated by integral membrane proteins, mechanotransduction dictated by the stiffness of the matrix, and the influence of surface chemistry and topography (reviewed by Guilakand colleagues^6^). However, the impact of niche geometry and its role in the regulation of the stem cell niche have not yet been fully explored. All epithelial tissues present common structural features, with stem cell niches generally identified as invaginations of the basement membrane^7^. During homeostasis and in the presence of wounding, stem cells divide and transient epithelial stem cells migrate unidirectionally whilst differentiating into mature, functional epithelial cell types.

In this study, we investigate how the geometry influences and controls the differentiation of stem cells, independently of imposed chemical and environmental conditions. As an exemplar, we focused on the limbal stem cell niche that serves the epithelium of the cornea, the transparent structure located at the front the eye. The limbus is a 1 mm-wide circular region separating the sclera from the cornea. It contains the palisades of Vogt, some of which are associated with limbal epithelial stem cell niches, called Limbal Epithelial Crypts (LECs). The presence, location and shape of the LEC were described for the first time by Dua *et al*.^8,9^. During the healing process and normal physiological turnover, human Limbal Epithelial Stem Cells (hLESCs) migrate towards the centre of the cornea and differentiate^10^ to constantly replace the corneal epithelial cells^11^. Such niche structures are common in all epithelial tissues and share certain characteristics including geometry, a stromal compartment known to influence epithelial cell phenotype through chemical signalling, and zonation in epithelial cell phenotype such that there is a stem cell, a progenitor or transiently amplifying cell zone, and a region where cells are terminally differentiated. This process is tightly regulated under the control of chemical signalling gradients^12^.

We hypothesise that in addition to the well-known cues associated with mechanical behaviour of the substrate and the physico-chemical gradients, geometry plays a key role in determining stem cell fate. To study this, we used two photon polymerisation (2PP) to create gelatin-based 3D crypts that mimic the precise geometry of the limbal stem cell niche. Multi-photon lithography is an advanced additive manufacturing technique capable of fabricating 3D micro/nano structures without a mask and with a possible feature size of below 100 nm. This is achieved through the absorption of multiple photons simultaneously, usually in the infrared range, and the requirements for such an event results in small region illumination with a sufficient intensity of photons to initiate polymerisation^13^. Following the creation of niche mimicking geometries using 2PP, we seeded hLESCs within the artificial niche and observed both proliferation and spatial variation in differentiation. We demonstrate statistically significant differences in the tendency to differentiate between cells located at the base of the niche and those situated towards the rim, supporting the hypothesis that stem cell fate is strongly influenced by their location within a niche and the geometrical details of where they reside.

Starting from histological sections described by Dua *et al*.^8^, crypt dimensions were identified and a tapered geometry was designed to replicate the niche by CAD. These structures were then fabricated by 2PP from Gelatin Methacrylate (GelMA) as U-shaped scaffolds with a diameter narrowing from 200 µm (rim) to 20 µm (base). GelMA presented a degree of methacrylation of 80.2% ± 6.1% according to NMR and this was confirmed by the TNBS assay (**Supplementary 1**). The scaffolds were successfully printed and well represented the CAD model, with a shape that narrowed at the base (**figure 1a-d**).

**Figure 1:**
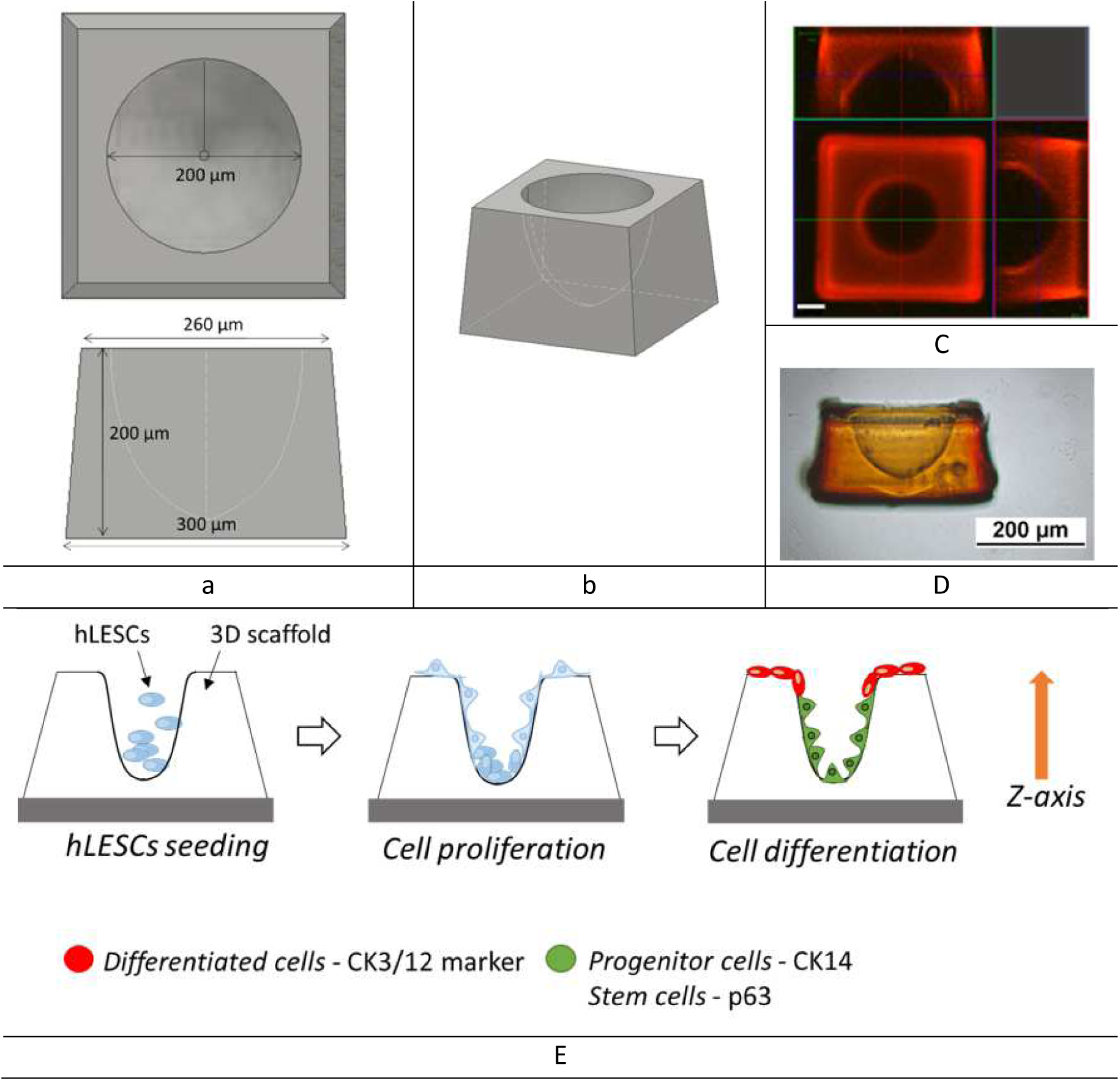
Limbal stem cell niche fabrication and scheme representing the assessment of stem cell response. (a) Top, side view and (b) 3D reconstruction of the CAD model developed to mimic the *in-vivo* structure; (c) Confocal images of top and side view of the 3D printed structure (scale bar 50 µm). (d) A brightfield image of the side view of the printed niche. (e) Graphical representation of the cell experiments: hLESCs are seeded into the printed scaffolds; cell proliferation is monitored and cell differentiation is assessed by CK3/12 (differentiation) or CK14 (limbal progenitor cell) marker expression.

The hLESCs were seeded inside the niches in xeno-free conditions to investigate their ability to repopulate the crypts and to study their differentiation. Commercially available hLESCs (CELLnTEC; Switzerland) displayed a typical polygonal epithelial morphology, and presented a homogenous monolayer of single-cells, without any colony formation (**Supplementary 2**).

After seeding, cells proliferated and were distributed along the length of the U-shaped scaffold, from the base to the top (**Supplementary 3**). Cell differentiation was studied along the z-axis and cytokeratin 14 (CK14) and cytokeratin 3/12 (CK3/12) were used as markers for limbal progenitor cells and differentiated corneal epithelial cells, respectively^14,15^. Zonation in marker expression was observed along the z-axis, presenting differentiated cells on the base (CK3/12 marker, labelled in green), and limbal progenitor cells on the top (CK14 marker, labelled in red) (**figure 2 and Supplementary 4**). As CK14 is a progenitor cell marker (rather than a stem cell marker, such as p63 as we report later), we conclude that the cells expressing this marker represent a population of cells at different stages of the differentiation pathway.

**Figure 2:**
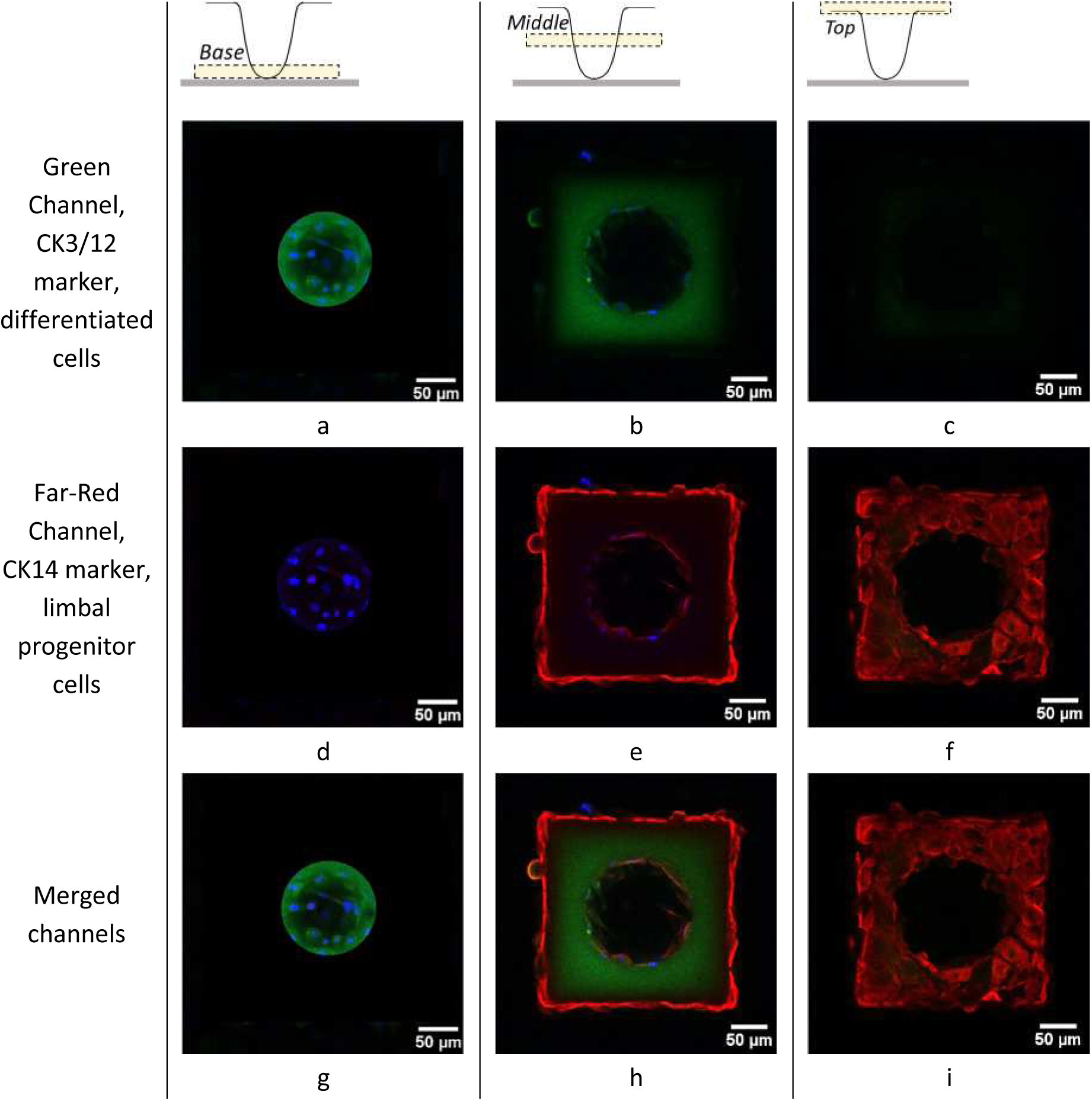
Zonation in marker expression was observed along the z-axis. Three z-positions representing (a,d,g) the base, (b,e,h) the middle, and (c,f,i) the top of the scaffold are reported. Cells were stained with CK3/12 (green; differentiated cells), CK14 (red; limbal progenitor cells) and Hoechst (blue; cell nuclei). (a-c) Green, (d-f) far-red and (g-i) merged channels are presented. All the channels are merged with Hoechst channel. The majority of the cells on the base expressed the differentiation marker, while at the top the majority expressed the stem cell marker. Images are representative of three independent experiments, performed with human Limbal Epithelial Stem Cells (hLESCs) obtained from CELLnTEC Advanced Cell Systems (P.2, single-donor).

Atomic force microscopy (AFM) was conducted to identify the stiffness in the z-direction, allowing the elastic modulus measurements to be performed at the same length scales at which cells interact with material^16^. Due to the length of the AFM tip, the base of the scaffold was not accessible, hence its properties could not be measured. For this reason, the base was measured on a flat scaffold, presenting the same printing parameters used for making the base of the U-shape scaffold. The results indicated that the printing process influenced the stiffness in the z-direction, however there are no studies so far that have shown correlation between stiffness and z-position of similar scaffolds. Future work will investigate this aspect in 2PP additive manufacturing, which may allow the production of biomaterials with stiffness gradients through the fabrication process. The top of the scaffold was stiffer than the base, presenting an average modulus of 54.85 ± 42.77 kPa and 14.33 ± 16.02 kPa, respectively (**figure 3**). The elastic modulus was in the same order of magnitude of the cornea also measured by AFM (7.5-110 kPa^17^). This range of values describes the local stiffness variations occurring in the transition from the limbal to the corneal tissue^18,19^. To understand if the z-position could influence cell differentiation, two flat scaffolds with different heights, 20 µm (Elastic modulus = 4.9 ± 2.5 kPa) and 200 µm (Elastic modulus = 28.7 ± 7.2 kPa), were produced, and hLESCs were seeded onto them. Immunostaining and confocal images were performed as before. The results indicated that the cells expressed predominantly the limbal progenitor cell marker (CK14) on both materials, independently of the height and stiffness (**Supplementary 5**). It is well known that the behaviour of cells is affected by the stiffness and the mechanical nature of the surfaces^20^, and also in the limbus there is some evidence that stiffness can impact on the stem cell differentiation^19^. However, in the data presented here, a difference in cell phenotype was not detected over the timeframe studied and warrants further investigation.

**Figure 3:**
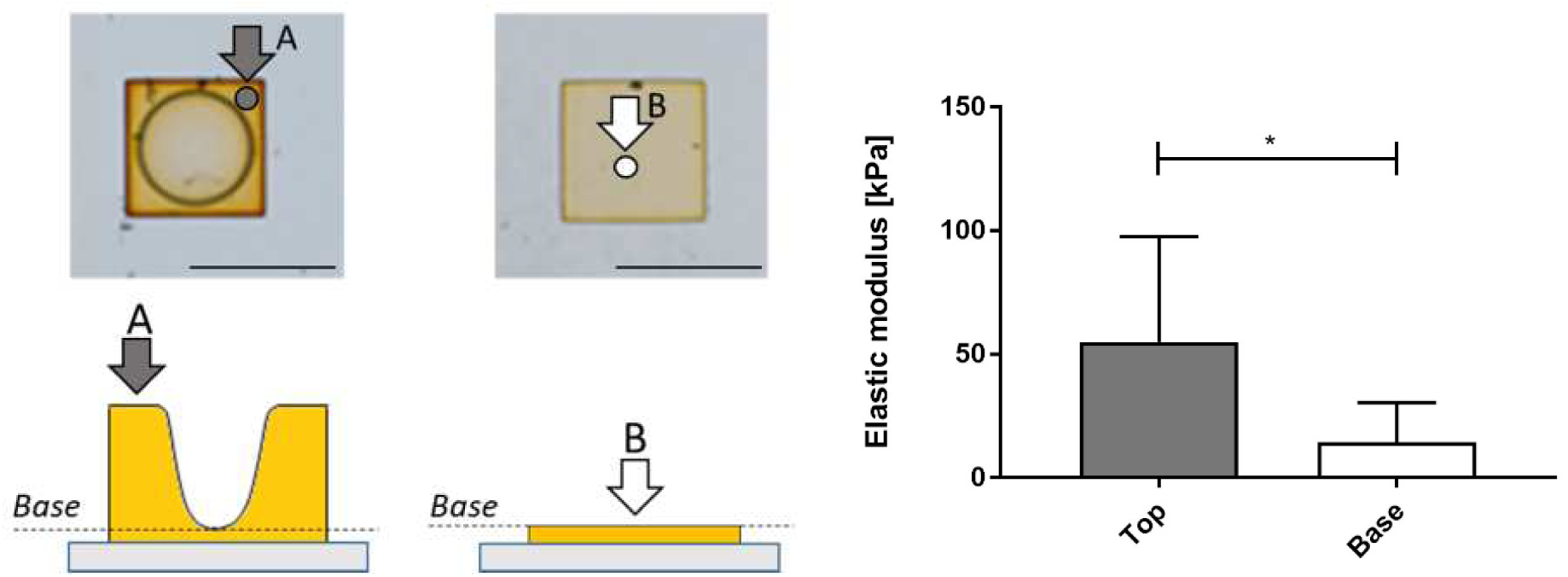
Surface elastic modulus of the scaffold according to the z-position, and representation of the areas on which the elastic modulus of the 2PP scaffold were measured by AFM. AFM indicated that the elastic modulus of the top (grey bar, kPa) is higher than the base (white bar, kPa) (n=3, p≤0.05). The arrows indicate where measures were taken.

As a further confirmation of the zonation in cell phenotype observed in the commercially available human primary stem cell line, we performed the same protocol by using freshly isolated cells from biopsy samples supported by an underlying feeder layer. Limbal epithelial cells were extracted from human cadaveric limbal biopsies and cultured using a feeder layer of irradiated 3T3-J2 fibroblasts and serum-supplemented media. The study of cell differentiation was conducted with CK3/12 as a differentiation marker, and p63 as a stem cell marker (**Supplementary 6**). The results also demonstrated a zonation in marker expression with cells at the base of the scaffold expressing the differentiation marker (CK3/12; green), while cells at the top expressed stem cell markers (p63; red) as seen with the commercially available cells. These results confirmed that the geometry itself is a decisive factor for the control of cell behaviour.

We postulate that the geometry created a milieu of cellular secretions inside the niche that enhanced cell differentiation at the base, and not at the top. Therefore, although we present here that geometry alone influences cell phenotype, the organisation of cells within the niche and indeed the control of their phenotype is influenced by other factors. Further studies to understand the interplay between geometry, chemical signalling and matrix stiffness is required to fully understand this phenomenon. Another parameter that could have had an influence in this process was the topography at the base of the scaffold. In previous articles it has been hypothesised that specific configurations, for example cells adhering on pillars separated by large distances (micro-patterns of 11 µm rather than 5 µm) can enhance the expression of late corneal keratinocyte markers^21,22^. A further study showed that cells seeded on restricted ECM contact triggered human epidermal stem cells to initiate terminal differentiation^23^.

Only a few other studies have attempted to replicate limbal niche structures ^24,25^ although these studies have not replicated the precise dimensions as described by Dua *et al*. ^8,9^ Ortega and colleagues did not illustrate zonation in phenotypic expression (using CK3 and p63 as markers of the differentiated and stem cell phenotypes respectively) when rabbit limbal epithelial cells populated ‘pockets’ which were 500 µm in height and 300 µm in diameter.^24^ In the paper by Levis and colleagues, their open ridge-like structures included human stromal fibroblasts and therefore the phenotypic response they reported in the limbal epithelial cell population cannot be attributed to geometry alone^25^. Overall, geometrical induced differentiation has not been achieved in any biologically relevant architecture which is the significant finding in this study. The results will have an implication in the design of novel scaffolds for the regeneration of epithelial tissues with the inclusion of stem cell niches.

In conclusion, we have demonstrated that geometry alone can influence epithelial stem cell fate, without the use of exogenous signalling molecules. We achieved this by producing niches with a replica *in vivo* geometry using additive manufacturing techniques that allow the geometrical freedom required to recreate structures at length scales appropriate for the production of stem cell niches. This method is flexible and can be applied to other epithelial tissues and as such, represents a significant advance in regenerative medicine. Uncovering of the underlying principles that govern cell response to niche geometry and stiffness, in addition to chemical signalling, will unblock the road to effective stem cell therapies and will ultimately lead to the emergence of new technological solutions in the form of implantable artificial niche based treatments. As such, precision engineering of these crypt structures holds potential to be a disruptive technology in the treatment of stem cell deficiencies.

## Supporting information

## Acknowledgments

This work was supported by the Engineering and Physical Sciences Research Council (EPSRC) and Medical Research Council (MRC) Centre for Doctoral Training (CDT) in Regenerative Medicine [EP/L015072/1], doctoral training grant awarded to E Prina. A short term scientific mission (STSM) awarded to E Prina was funded by the Biomedicine and Molecular Biosciences EU COST Action BM1302 ‘Joining Forces in Corneal Regeneration Research’. The two photon polymerisation fabrication was supported by the EPSRC the grants Multifunctional Additive Manufacturing [EP/K005138/1] and the EPSRC Centre for Innovative Manufacturing in Additive Manufacturing [EP/I033335/2]. We also wish to acknowledge Robert Markus (University of Nottingham) for assistance with confocal microscopy with equipment funded by the Biotechnology and Biological Sciences Research Council (BBSRC) [BB/L013827/1] and Xinyong Chen (University of Nottingham) for assistance with atomic force microscopy.

## Author Contributions

The manuscript was written through contributions of all authors. Elisabetta Prina carried out the majority of the experimental work and manuscript preparation for this paper. Expertise in limbal stem cell isolation, culture and characterisation was provided by Laura Sidney, Marina Bertolin, and Stefano Ferrari. Two photon amenable photoinitiators were synthesized by Maximilian Tromeyer, Robert Liska and Jonathan Moore. Insights into corneal histology were provided by Andrew Hopkinson and Harminder Dua. The work was conceived, organised and made possible with funding awarded to Jing Yang, Ricky Wildman and Felicity Rose. Manuscript drafting and submission was led by Ricky Wildman and Felicity Rose. All authors have given approval to the final version of the manuscript.

## Materials and methods

### GelMA synthesis

Gelatin Methacrylate (GelMA) was synthesized as described by Nichol *et al*.^26^. Porcine skin gelatin (10% (w/v) Type A, 300 bloom, from Sigma-Aldrich, UK, G2500) was dissolved in Phosphate Buffered Saline (PBS, Gibco, Thermo Fischer Scientific, UK, BR014G) at 80°C and stirred until fully dissolved. Methacrylic anhydride (8% (v/v) Sigma-Aldrich, UK, 276685) was added at 0.5 mL/min and stirred at 60°C for 3 hours. To quench the reaction, the solution was diluted five times with PBS and stirred for a further 30 minutes. The solution was dialyzed against distilled water using 8 kDa cut-off dialysis tubing (BioDesignDialysis Tubing™ D118, USA) for 1 week at 50°C. The dialysis water was changed 3 times per day to eliminate the salt and the methacrylic acid. The solution was filtered through a 40 µm filter and lyophilized (Edwards Modulyo freeze dryer, IMA Edwards, UK) for 3 days. The freeze-dried GelMA was stored at −20°C until use.

### Degree of crosslinking: NMR and TNBS assay

The percentage of methacrylation was quantified using ^1^H Nuclear Magnetic Resonance at 400 MHz (NMR, DPX UltraShield 400MHz, Bruker UK ltd, UK). The spectrum was collected at room temperature (RT). Phase and baseline corrections were applied before obtaining the integrals of the peaks of interest. MestReNova software (v6.0.2) was used to analyse the spectra (64 scans). Freeze dried GelMA and gelatin were dissolved in Deuterium Oxide (D_2_0, Sigma-Aldrich, UK, 151882) at a concentration of 10 mg/mL. The presence of the double bond signal (5.35-5.60 ppm) indicates the addition of the methacrylate vinyl group. The percentage of methacrylation was calculated using the method described by Hoch et al.^27^. Briefly, the integral of the methylene peak (d = 2.7 ppm to d = 2.9 ppm) was used for the quantification of the lysine signal. The degree of substitution was determined by comparing the lysine signals of the modified and unmodified gelatin according to the equation 2.1. The intensity of the aromatic region (7-7.5 ppm) was used as a reference.

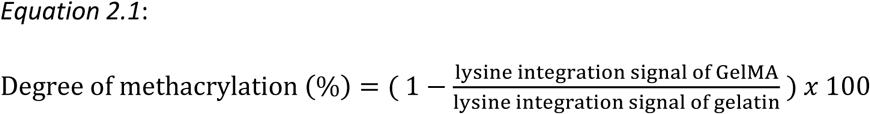

The degree of methacrylation was also confirmed by using the 2,4,6-Trinitrobenzene Sulfonic Acid (TNBS, Sigma-Aldrich, UK, P2297) assay. Gelatin and GelMA (4 mg), were dissolved in 0.25% (*v/v*) of TNBS (diluted in 0.01 M sodium hydrogen carbonate solution, Sigma-Aldrich, UK, 367176), and incubated at 40°C for 3 hours. Hydrochloric acid (3 mL, 6N, Alfa Aesar, UK, 44921) was added and the solution incubated at 80°C for one hour to dissolve any insoluble matter. After cooling down to room temperature, 4 mL of deionized water (dH_2_O) was added and the absorbance was measured at 345 nm using a spectrofluorometer (Tecan Infinite^®^ M200 microplate reader, UK). The absorbance value indicated the presence of free amino groups, and the degree of crosslinking calculated according to the equation 2.2:

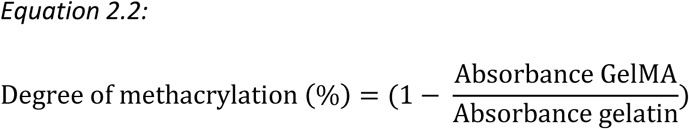

#### Two photon polymerization of GelMA to create crypt structures

The polymeric solution was prepared by mixing GelMA foam to the photoinitiator P2CK. GelMA (15% *w/v*) and the photoinitiator P2CK were investigated in order to obtain a suitable formulation for the current application. The photoinitiator, P2CK, was prepared according a published protocol^28^ (Vienna University of Technology) or by a modified procedure (University of Nottingham). GelMA (15% *w/v*) was dissolved in PBS, and the photoinitiator P2CK was added to obtain a final concentration of 0.3% *w/v*. To fabricate micro-structures replicating the dimensions of the limbal stem cell niche, a two photon polymerization (2PP) system (NanoScribe Photonic Professional, GmbH, Germany) with a femtosecond laser emitting at 780 nm in galvo scan mode was used. The structure was developed using CAD 3D Autodesk Inventor software and the printing parameters were set by the Describe software. The laser beam was focused using a conventional 63x microscope objective (Zeiss, NA 1.4, with oil immersion media), which was mounted on a linear stage for vertical positioning. The laser writing system was controlled by the Nanowrite software. A drop of each individual gel was cast between a 22×22 mm glass slide (thickness n. 1.5), and an 18×18 mm cover glass (thickness n. 1.5), separated by a 300 µm spacer (figure 2.1). The printing process was obtained by tuning laser power, scan speed and the hatching distance (distance between each layer). After printing, the structure was ‘developed’ by immersing the sample in PBS at 40°C for 1 hour to remove the non-polymerized hydrogel; the sample was then left at 4°C overnight. Irgacure 2959 solution (0.1% w/v) was added and the scaffold UV cured (365 nm long wave UV lamp at 15W) for 10 minutes to increase the stability of the structures.

**Figure 2.1:**
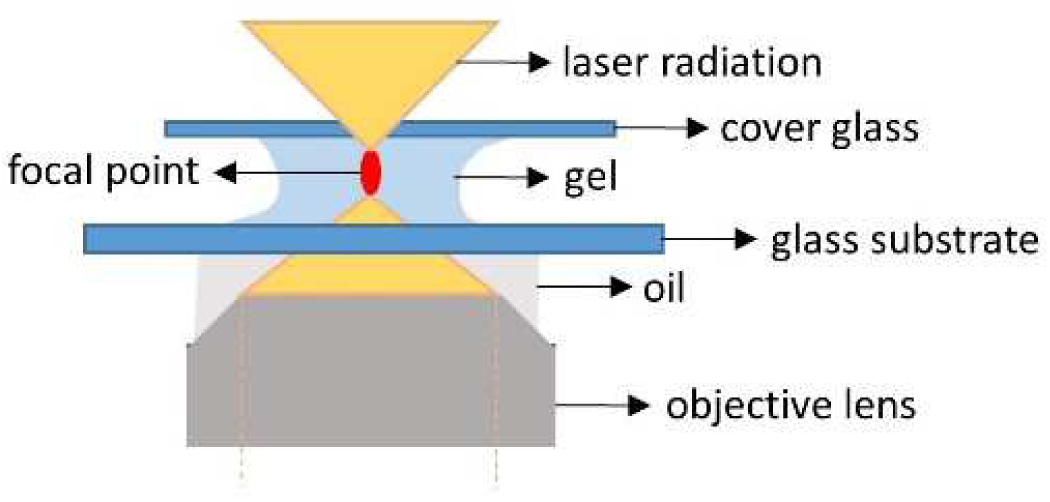
Configuration of gel set up within the 2PP system. The gel is sandwiched between two sheets of glass and the laser is focused with 25x or 63x objectives.

### Determining the surface elastic modulus of the gel

To measure the elastic modulus of the surface of the gel, AFM (MFP-1D instrument, Asylum Research, USA) was performed. A thin cantilever (nominal spring constant between 0.070 N/m and 0.025 N/m) with a silicon nitride tip (MSNL-10 type, probe A, from Bruker Nano, Coventry, UK) was used. The cantilever was positioned on the area at top of the scaffold, and on a printed flat scaffold (height=20µm) to emulate the base of the scaffold. The force graph was recorded, and the elastic modulus (E) calculated from the approaching force curves based on the Hertz model^29^, according to the equation:

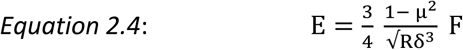

where F is the applied force, R is the radius of the probe, δ is the indentation od the samples, E is the elastic modulus and µ is the Poisson’s ratio. Before starting the test, tip sensitivity and the calculation of the spring constant was conducted. Three experiments were conducted for each condition (n=3). The measurements of each experiment were grouped and a two tails unpaired t-test was conducted in Prism 6 (GraphPad Software, v6.01). Difference was considered statistically significant when the p-value was lower than 0.05 (p≤0.05).

### Cell culture

#### Xeno-free human limbal stem cells

Commercial human Limbal Epithelial Stem Cells (hLESCs) were obtained from CELLnTEC Advanced Cell Systems (P.2, single-donor, Switzerland) and cultured in serum free media (CnT-Prime, Epithelial Culture Medium, CnT-PR, CELLnTEC Advanced Cell Systems, Switzerland) with the addition of 100 units/mL of penicillin, 0.1 mg/mL of streptomycin and 0.25 μg/mL of amphotericin B and kept at 37°C, 5% CO_2_. Cells were cultured in 75cm^2^ culture flasks, at a cell density of 4,000 cells/cm^2^.

### Primary cells extracted from human corneal biopsies

Tissue-derived human Limbal Epithelial Cells (td_hLESCs) were collected from scleral-corneal tissue (obtained with written consent from the next of kin to be used for research, according to the directives set by the Italian Centro Regionale Trapianti and Centro Nazionale Trapianti and only when tissues were not suitable for transplantation) and preserved in media before processing. The biopsy was composed of cornea, limbus and conjunctiva which was dissected into quarters, then the cornea and the conjunctiva were removed. The limbus was fragmented and soaked in PBS. Trypsin/EDTA (0.05% / 0.01% *w/v*; ThermoFisher, Italy, 25300-62 was added (10 mL) to digest the tissue for 30 min at 37°C. After 30 min, media containing fetal bovine serum (FBS) was added to inhibit the trypsin action. The solution was centrifuged for 5 min at 1000 rpm (Thermo Scientific-Megafuge 1.0). After removing the supernatant, the pellet was re-suspended in fresh media and the cells were counted. This cycle was repeated three times to extract the majority of the cells. Cells were plated onto a 24 well plate (30,000 cells/well), in co-culture with pretreated with irradiated 3T3-J2 cells. The standard control medium consisted of Dulbecco’s Modified Eagle Medium (DMEM, ThermoFisher, Italy, 21969) and Ham’s F12 (F12) (DMEM/F12 2:1, Ham’s F-12, ThermoFisher, Italy, 21765) supplemented with 10% (*v/v*) FBS (ThermoFisher, Italy, 0101-145), 50 μg/mL penicillin-streptomycin (P/S, Euroclone, Italy, ECB3001D), 4 mM glutamine (Euroclone, Italy, ECB3000D), 5 μg/mL insulin (Humulin R, Lilly, Canada, HI0210), 0.4 μg/mL hydrocortisone (Flebocortid Richter, Sanofi, Italy, AIC 013986029), 0.18 mM adenine (Adenine grade I, Pharma Waldhof GMBH, Germany, 4010212), 8.1 μg/mL cholera toxin (Cholera Toxin QD, List Biological Laboratories, USA, C8052-1MG), 2 nM triiodothyronine (Liotir, IBSA, Italy, AIC 036906016) and 10 ng/mL epidermal growth factor (EGF, GMP Cellgro, Cell Genix GmbH, Germany, 1016-050).

### Feeder layer preparation for the cultivation of td-LECs

Td_LESCs were cultured on irradiated 3T3-J2 cells, used as a feeder layer. 3T3-J2 cell media consisted of DMEM supplemented with 10% (*v/v*) foetal calf serum (ThermoFisher, Italy, 03990017M), 50 μg/mL penicillin-streptomycin (P/S, Euroclone, Italy, ECB3001D), and 4 mM glutamine (Euroclone, Italy, ECB3000D). Once confluent, the cells were trypsinized and centrifuged. After removing the supernatant, the cells were counted and placed in a 50 mL tube with a maximum of 20 mL cell culture media and 1 x 10^6^ cells/mL and irradiated with a final dose of 60 Grays (Gy) (X-Ray Irradiator, Faxitron-Cell Rad, 2328A50142). To prepare the feeder layer for the extracted limbal stem cells, 80,000 cells/well were plated in a 24 well plate.

#### Immunocytochemistry to determine cell phenotype

To study limbal cell marker expression, cells were dual-stained with CK14 and P63 (limbal progenitor and stem cell markers respectively) and CK3/12 (differentiated corneal epithelial cell marker). After fixation, cell membrane permeabilisation was carried out in 0.1% v/v triton X-100 (Sigma-Aldrich, UK, T8787) in PBS for 10 min at RT followed by washing with PBS three times for 5 min. Blocking solution was composed of PBS containing 1% (*w/v*) of Bovine Serum Albumin (BSA, BDH Prolabo^®^, VWR, USA, 421501J), 0.3 M glycine (Sigma-Aldrich, UK, G7126), and 3% (*v/v*) of serum from the animal in which the secondary antibody was raised (goat serum, Sigma-Aldrich, G9023); samples were exposed to blocking solution for one hour at RT. Primary and secondary antibodies were diluted in wash buffer (1% (*w/v*) BSA, 0.3M glycine in PBS; Table 2.1). Samples were incubated in the primary antibody overnight at 4°C followed by washing in buffer and the secondary antibodies added for one hour at RT. Samples were washed three times for 5 min in PBS and nuclei stained with Hoechst (B2883, Sigma-Aldrich, UK) for 10 min at RT. The Hoescht dye was removed, samples washed with PBS and mounting media added (Fluorescent mounting medium Dako, VWR, UK, S3023) prior to analysis by confocal microscopy.

**Table 2:**
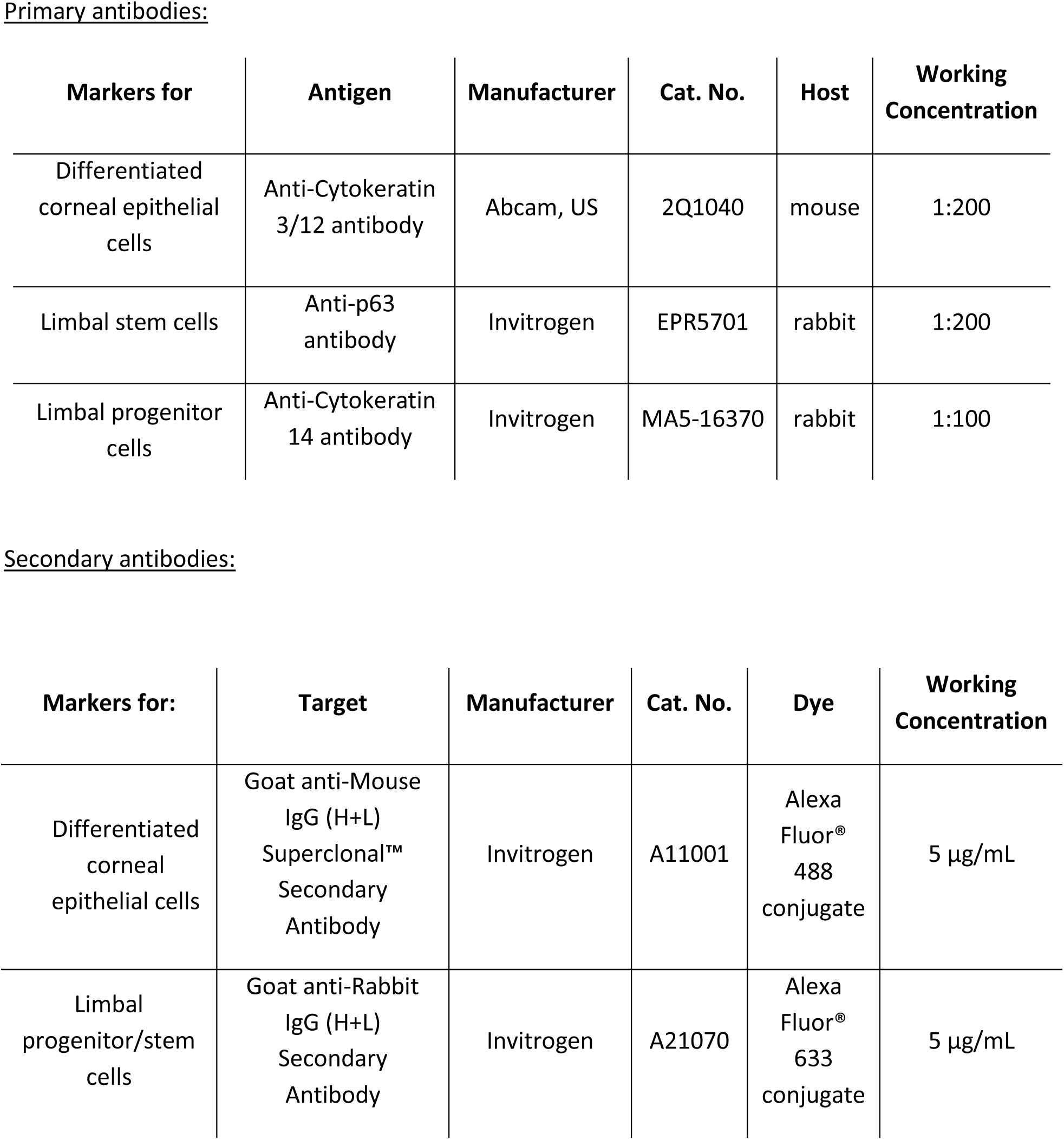
Primary and secondary antibodies used in immunofluorescence.

