## Supplementary material for "Geometry alone influences stem cell differentiation in a precision 3D printed stem cell niche"

**Supplementary 1: Degree of GelMA substitution**

The degree of substitution (DS) calculated with NMR was 80.2% ± 6.1 (figure S.1). These results were confirmed using the TNBS assay, where the average of three batches was 83.5% ± 4.9. In table S.1, the results of each batch are reported.

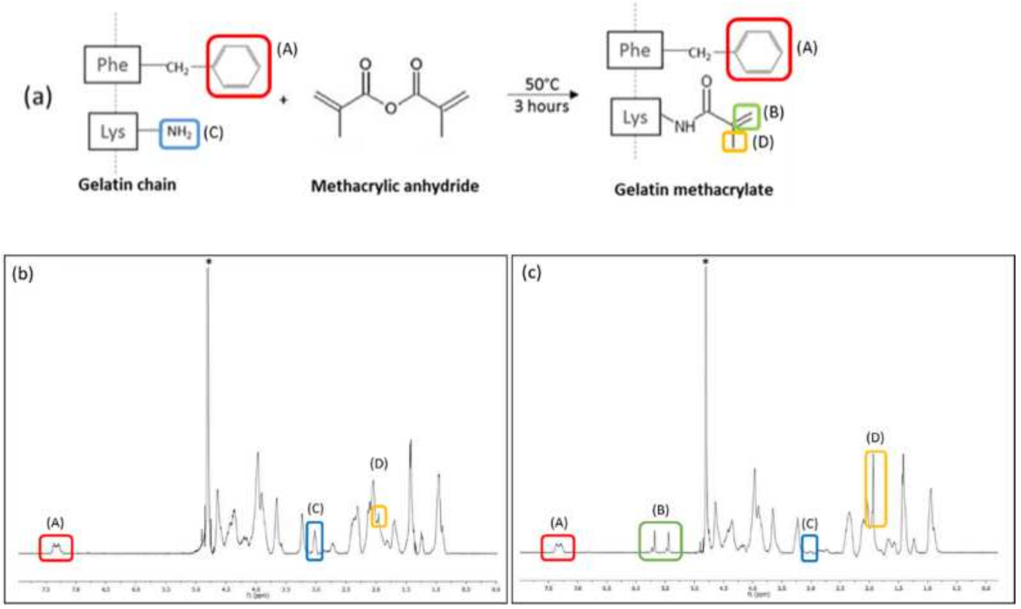


**Figure S.1: Gelatin methacrylate characterization.** (a) Schematic representation of methacrylate substitution of the primary amines of gelatin. NMR spectra of 10 mg/ml of (b) gelatin and (c) GelMA recorded in $D_20$ (*) at RT. Both graphs present the peak at 7.2 ppm associated with the aromatic groups (peak A, red). In the GelMA spectrum, the lysine signal (peak C, blue) at 2.9 ppm decreases while the methacrylate vinyl group signal at d = 5.4 ppm and d = 5.7 ppm (peak B, green) and the methyl group signal at 1.8 ppm increase (peak D, yellow), indicating that the reaction occurred.

**Table S.1: DS of three different batches of GelMA measured by TNBS assay.** Average ± SD (n=3).

| | ***Batch 1*** | ***Batch 2*** | ***Batch 3*** | ***Average ± SD*** |
|---|---|---|---|---|
| ***Degree of substitution*** | 84.2 | 78.3 | 88.0 | 83.5 ± 4.9 |

**Supplementary 2: Cell culture systems**

To study the behaviour of limbal stem cells seeded on the 3D printed niches, two protocols were used. Firstly, commercial human Limbal Epithelial Stem Cells (hLESCs, passage 2), CELLnTEC (Switzerland) were harvested and cultivated by using the basal media CnT-Prime, an animal-component-free culture medium developed for the isolation and expansion of epithelial cells. In the second protocol, tissue derived human Limbal Epithelial Cells (td_hLESCs) were extracted directly from the limbus of scleral-corneal tissues (obtained with written consent from the next of kin to be used for research, according to the directives set by the Italian Centro Regionale Trapianti and Centro Nazionale Trapianti and only when tissues were not suitable for transplantation), and seeded on a feeder layer of irradiated 3T3-J2 with serum containing-media. Both systems were able to support cell growth which displayed a typical polygonal epithelial morphology. In figure S.2, representative cell morphologies are shown. The method used in the treatment of LSCD is based on the expansion of *ex-vivo* LESCs on inactivated lethally irradiated 3T3-J2 fibroblasts as a feeder layer. However, translational research aims to reduce the risk of xenotoxicity and more effort has since been made to develop animal-free culture systems[30].

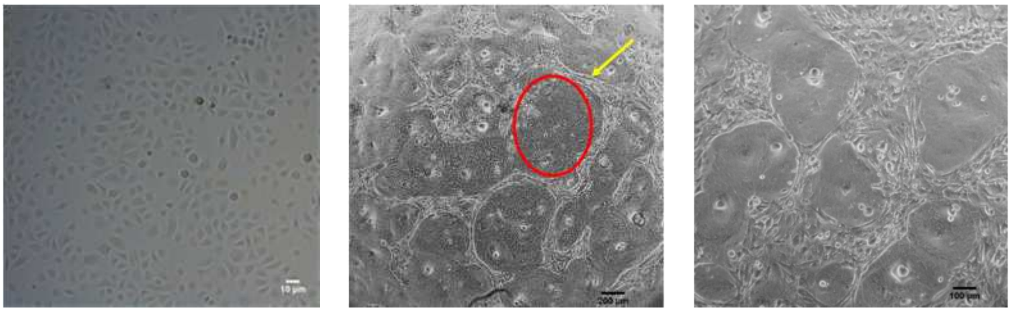

a b c

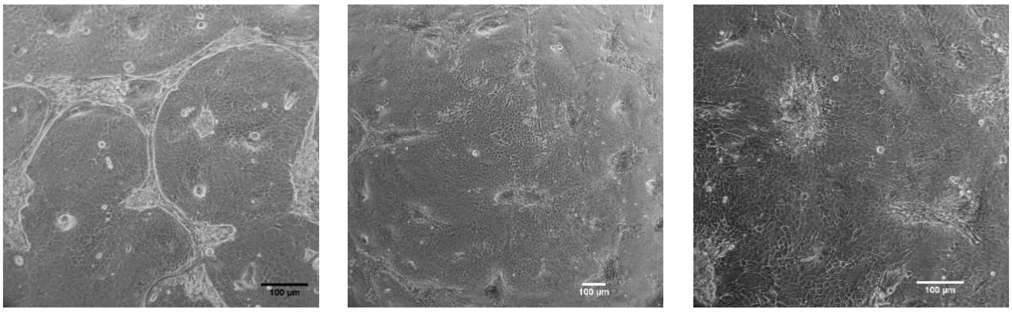

d e f

**Figure S.2: Cultivation of human limbal epithelial cells (hLESCs).** (a) Commercially available human limbal epithelial cells (hLESCs) were cultured in serum-free and feeder layer-free medium. They did not display colony formation. (b-f) Cells extracted directly from a limbal biopsy (td_hLESCs) were seeded on an irradiated 3T3-J2 layer. (b-d) The epithelial cells formed colonies (red circle), pushing away the fibroblasts (yellow arrow); (e-f) the epithelial cells became confluent, and the remaining feeder layer covered a small area, and were removed when changing the medium.

<u>**Supplementary 3: Limbal stem cells were distributed along the niche on the printed structures**</u>

A suspension of the commercially available hLESCs or a feeder layer of irradiated 3T3-J2 cells and td_hLESCs were seeded inside the scaffold, and their distribution along the z-axis was monitored by Hoechst 33258 staining. For both cell culture methods, cells were distributed along the scaffold, from the base to the top (**figure S.3**).

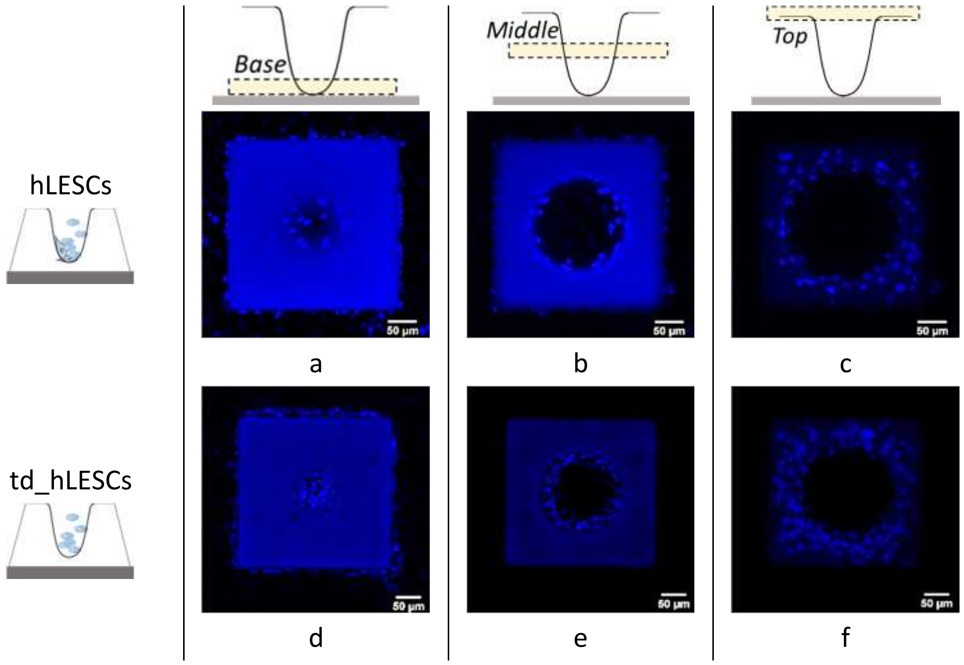


**Figure S.3: Distribution of primary cells along the z-axis**. (a-f) z-stack confocal images of Hoechst-stained cells at the base, middle and top of the scaffold. (a-c) hLESCs and (d-f) td_hLESCs are distributed along the z-axis of the scaffolds.

**<u>Supplementary 4: zonal expression of hLESC, progenitor and differentiated cell phenotype according to the z-position</u>**

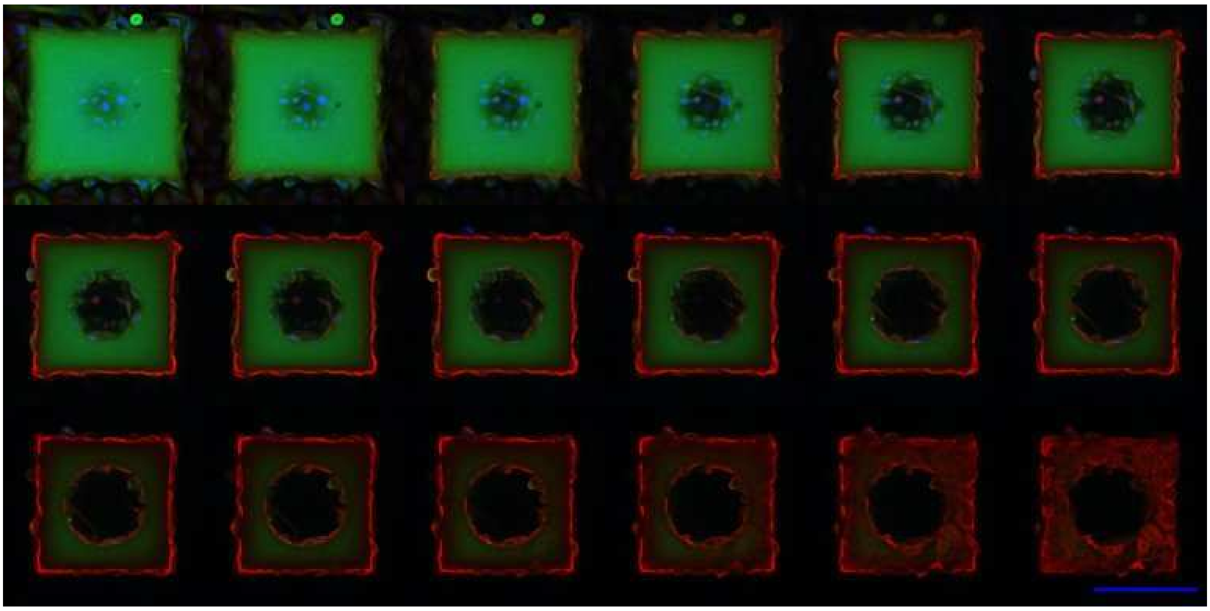

**Figure S.4**: Montage of the confocal images in z-stack. HLESCs were stained with CK3/12 (green; differentiated cells), CK14 (red; limbal progenitor cells) and Hoechst (blue; cell nuclei), and all channels were merged. Each image represents a z-plane of the scaffold moving from the base (top-left) to the top of the scaffold (bottom-right). Scale bar = 200 µm.

### Supplementary 5: hLESC expression according to the stiffness

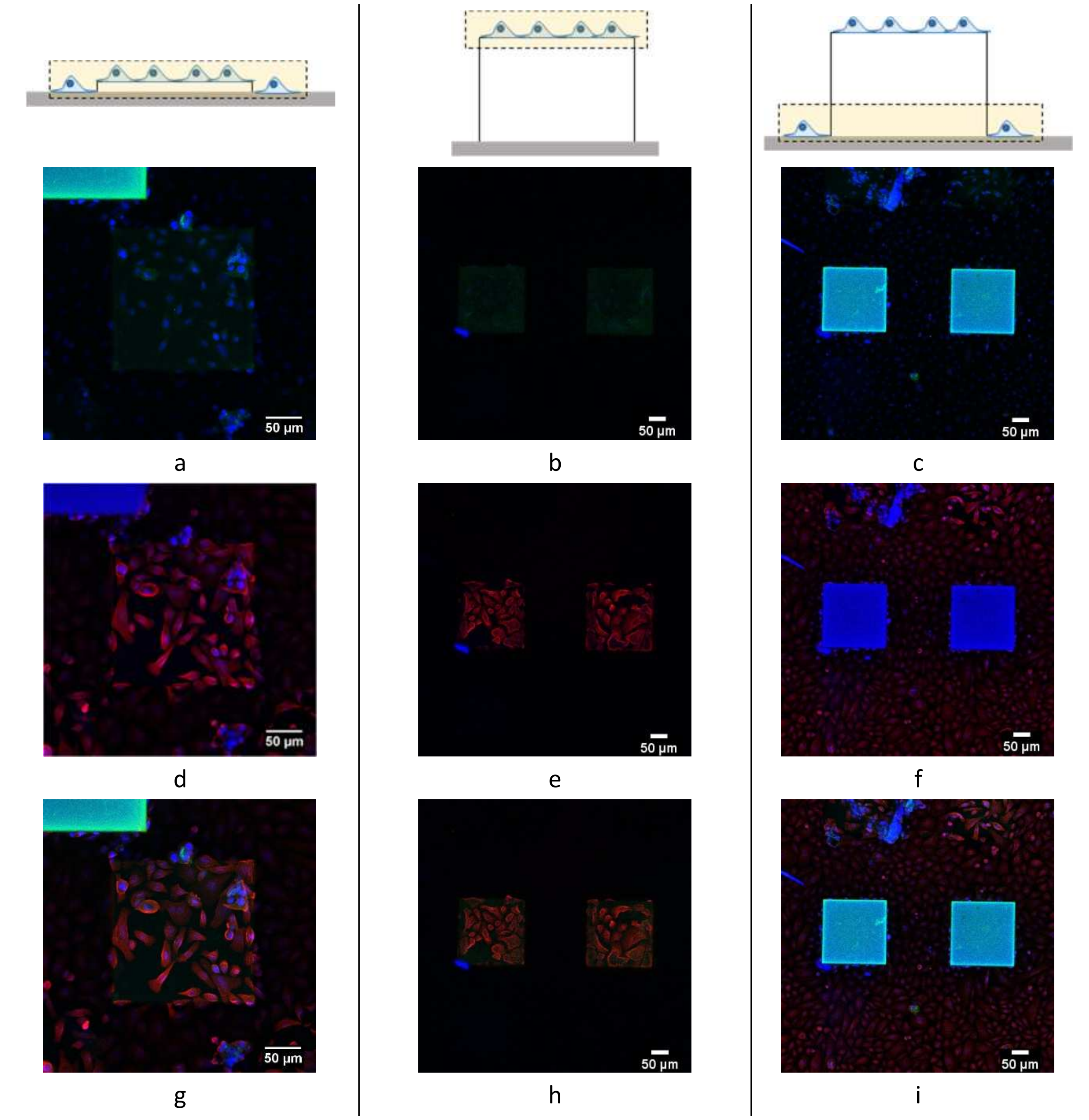


**Figure S.5: Confocal images of hLESCs on flat surfaces stained with CK3/12 (green), CK14 (red) and Hoechst (blue).** Cells were seeded on (a,d,g) flat scaffolds of 20 µm and (b,e,h) 200 µm height, and on (c,f,i) the glass surface. Green, far-red and merged channels are presented in the first (a-c), second (d-f) and third row (g-i), respectively. All the cells in these three conditions (N=3) expressed predominantly the CK14 limbal progenitor cell marker (red).

**Supplementary 6: td_hLESCs differentiated according to z-position**

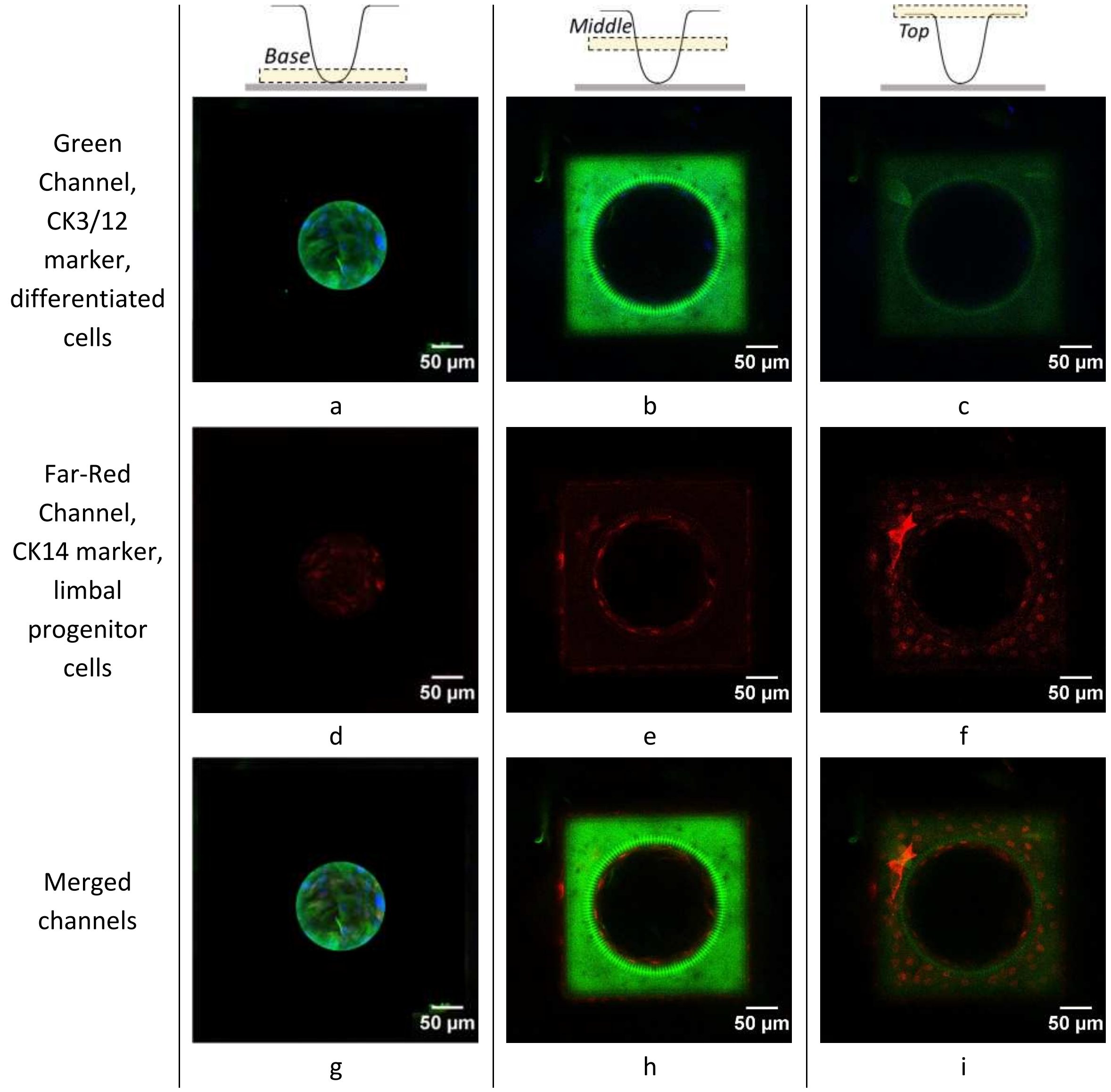


**Figure S.6: Series of confocal Z-stack images showing td_hLESC seeded within the scaffolds.** Three z-positions representing the base (a,d,g), the middle (b,e,h) and the top (c,f,i) of the scaffold are reported. Cells are stained with CK3/12 (green), CK14 (far-red) and Hoechst (blue). Green, far-red and merged channels are presented in the first (a-c), second (d-f) and third row (g-i), respectively. The majority of the cells on the base expressed the differentiation marker (CK3/12; green), while on the top they expressed the limbal progenitor cell marker (CK14; red).
